# Constrution of a genetic map of rils derived from wheat (*T. aestivum* L.) varieties pamyatii azieva x paragon using high-throughput snp genotyping platform kasp - *competative allele specific PCR*

**DOI:** 10.1101/740324

**Authors:** K Yermekbayev, S Griffiths, M Chettry, M Liverington-Waite, S Orford, A Amalova, S Abugalieva, Y Turuspekov

## Abstract

The main purposes of the study were i) to develop a first mapping population for bread wheat grown in Kazakhstan, ii) to construct its genetic map for further identification of genes associated with important agronomic traits.

To the best of our knowledge this is the first segregating population and genetic map developed for Kazakh bread wheat. The work is an example of how plant breeding programs in Kazakhstan have started successfully deploying next generation plant breeding methods.

The KASP (Compatative Allele Specific PCR) technology of LGC Group and SNP DNA-markers have been exploited to genotype and build a genetic map of the segregating population. The total length of the map was 1376 cM. A total 157 out of initial 178 SNP markers used formed 26 linkage groups leaving 1 duplicated and 20 unassigned markers. The threshold distance between markers was set ≤ 30 cM. Therefore, two linkage groups were obtained for chromosomes such as 2A, 2B, 2D, 3A, 5A, 6B and 7A. Despite one duplicated and 20 unassigned markers, the 157 KASP SNP markers that were mapped spanned A, B and D genomes of wheat. Kosambi Mapping function was employed to calculate recombination units between makers. RILs were developed through SSD method up to F4 generation. Almost 97% of identified alleles were useful in evaluating the population’s genetic diversity; the remaining 3% showed no outcome. As a result, 77 DNA markers were mapped for A, 74 for B and 27 for D genomes. The mapping population will be genotyped using high marker density array planform such as Illumina iSelect to obtain a genetic map with a relatively high coverage. Then, the population and high-resolution genetic map will be used to identify genes influencing wheat adaptation in Kazakhstan.

## Introduction

Wheat is a cereal crop. In spite of the fact that there are many slightly differing theories of its origin, the Fertile Crescent is mainly considered to be the place where wheat originated and was domesticated (CW 1995) (Nesbitt and Samuel 1998) (Gustafson, Raskina et al. 2009). The three genomes (AABBDD) of hexaploid wheat (*Triticum aestivum* L. ssp. *aestevium*) derived from diploid (DD) *Aegilops tauschii* and tetraploid (BBAA) wild emmer (*Triticum turgidum* L. (Tell), ssp. *dicoccoides*) which in turn inherited its AA genome from diploid *Triticum urartu* and BB from either diploid *Aegilops speltoides* or some other extinct species (Wang, Luo et al. 2013) (Feldman, Liu et al. 1997). Since its cultivation bread wheat has been a crop of great importance which is currently supplying 20% of the total world population’s calories (Bushuk 1997). Therefore, hexaploid (2n=6x=42) bread wheat with the most complex genome and origin of history is a vital food and feed crop in the world. Its importance is particularly high in Kazakhstan, because i) most of the local dietary commodities consisted of wheat flour (Eken, Spanbayev et al. 2016) ii) Kazakhstan is one of the three giant world wheat exporters, currently ranked as one of the four major wheat exporters which are collectively known as the world’s “bread basket” countries along with Ukraine and Russia (Swinnen, Burkitbayeva et al. 2017).

Although agriculture is one of the key sectors of the national economy of Kazakhstan, a literature review showed a shortage of genomic experiments in the country’s current cereals breeding programs. A paucity of well-trained research staff, costly reagents and lack of sufficient funding and facilities could be the main reason why such fundamental and pre-breeding project proposals have not been carried out so far. Nonetheless, with an initiative of Prof Y. Turuspekov and Prof S. Abugalieva (Institute of Plant Biology and Biotechnology, Almaty) the genetic diversity of commercial/elite cultivars of wheat (Abugalieva, Ledovskoy et al. 2010) (Abugalieva, Volkova et al. 2012, Turuspekov, Plieske et al. 2017) (Turuspekov, Baibulatova et al. 2017), barley (Turuspekov, Ormanbekova et al. 2016) (Genievskaya, Almerekova et al. 2018), and oat (Abugalieva, Sereda et al. 2013) (Ashimova, Yermekbaev et al. 2016) have been evaluated using different types of molecular markers. Importantly, genes associated with essential agronomic traits have been identified through the exploitation of these new tools and resources. However, there is still a wealth of genetic information that has not been properly captured to intensify breeding programmes yet, because of either disconnection between plant geneticists and breeders or a low level of motivation for plant breeders to use molecular genetics approaches. Thus, our next aim is to carry out pre-breeding programs employing quantitative genetics approaches and Marker Assisted Selection. As animal and plant breeding is mainly based on conventional methods in developing countries, like Kazakhstan, our main objective is to carry out systematic plant breeding by coupling conventional plant breeding methods with modern genomic selection approaches. The reason for targeting developing countries is that the most of them possess enormous agronomic potential to sustain future food security of the world.

## Materials and methods

### Development of segregating/mapping population

A segregating population of Recombinant Inbred Lines (RILs), was developed from a cross between the spring wheat cultivars, Pamyati Azieva (Pam) and Paragon (Par), through Single Seed Descent (SSD) bulk method (Tee and Qualset 1975) which is widely used among breeders for the last three decades (Fischer and Rebetzke 2018). RILs are a powerful tool for the mapping of genes (Broman 2005) and have been used for the identification and validation of QTL (quantitative trait loci) underpinning central agronomic traits of an important staple cereals such as wheat (Ren, Wang et al. 2018), barley (Swarcewicz, Sawikowska et al. 2017), rice (Ren, Wang et al. 2018), chickpea (JUKTE 2018), soybean (Grunvald, Torres et al. 2018) and maize (Pan, Xu et al. 2017). The first parent, which is well adapted to Kazakhstan - Pamyati Azieva, used in the cross, originated from Sibirskii NIISKH in Omsk, Russia, and is widely grown in Kazakhstan because it was included into the recommended list of the State Register of the country. Paragon is a UK spring wheat variety. It has been used extensively in the UK’s wheat pre-breeding programs as a parent for the development of Nested Association Mapping (NAM) populations, Near Isogenic Lines (NILs), and mutant populations (Wingen, West et al. 2017) because it was recognised as the bread making quality benchmark in the UK. Once, these two elite varieties were crossed and F1 hybrid lines obtained, in each round of the bulk one seed was taken to develop next generation, that is one seed from F2 was used to obtain F3. Only one seed of F4, meanwhile, was employed to form F4 hybrids. The F4 generation was used for genotyping and map construction. All hybridization experiments and development of RILs carried out in glass houses and Controlled Environment Rooms (CER) of John Innes Centre (JIC), UK, between 2012-2014.

### DNA extraction from Pam x Par RILs and DNA preparation for genotyping using KASP

In order to extract DNA from Recombinant Inbred Lines, five seeds of each line were sown into Petri dishes and put for 2 days at 4°C for stratification. After that, they were transferred to an incubator with temperature rages of 27-30°C for 5-6 days for germination. Dellaporta methodology (Dellaporta, Wood et al. 1983), with changes, was used to isolate DNA from 5-6 days old wheat seedlings. Extracted genomic DNA has been subjected to further purification using Qiagene Mini spin columns. Quality and quantity (A260/A280) of extracted DNA were evaluated in Nanodrop spectrophotometer with further visualization on 1% agarose gels to assess integrity. DNA of each RILwas standardized to a concentration of 10 ng/μl for genotyping using the KASP platform. The main priority of KASP genotyping platform over old systems is that it does not require much time for PCR product visualization. Therefore, it enormously alleviated an old time-consuming visualization method, gel-based electrophoresis, which uses mutagenic wet chemicals such as ethidium bromide. In addition, using KASP technology the unique insertions, deletions and single base (one nucleotide) changes in the genomes can also be detected which makes it powerful tool to replace old tedious method.

### Genetic map construction

Initially, more than three hundred SNP DNA markers designed by Bristol University (the information about DNA markers can be obtained here www.cerealsdb.uk.net (Wilkinson, Winfield et al. 2012)) were used to genotype Pamyati Azieva and Paragon. This whether those molecular markers were polymorphic or not. Once polymorphic markers were identified which distinguished parents genetically, they have been employed to genotype the RIL population. To construct genetic linkage map of RILs, MapDisto software was used initially (http://mapdisto.free.fr/) (Heffelfinger, Fragoso et al. 2017). However, the genetic map distance was re-estimated using an omnibus statistical software environment R. The reason is that R is flexsible tool to conduct statistical calculations for the dataset to improve a genetic map quality. In both cases Kosambi mapping function (Kosambi 2016) was used to calculate recombination fractions and then, convert units to centiMorgans. Logarithm of odds (LOD score) was not lower than three. In total 72 SNP molecular markers have been mapped for A, 67 for B and only 19 for D genomes showing importance of development of more polymorphic markers for D genome to deploy its genetic potential for marker-trait association analysis. No markers have been mapped for 3D and 4D chromosome of wheat (Table 1).

**Table 1.**
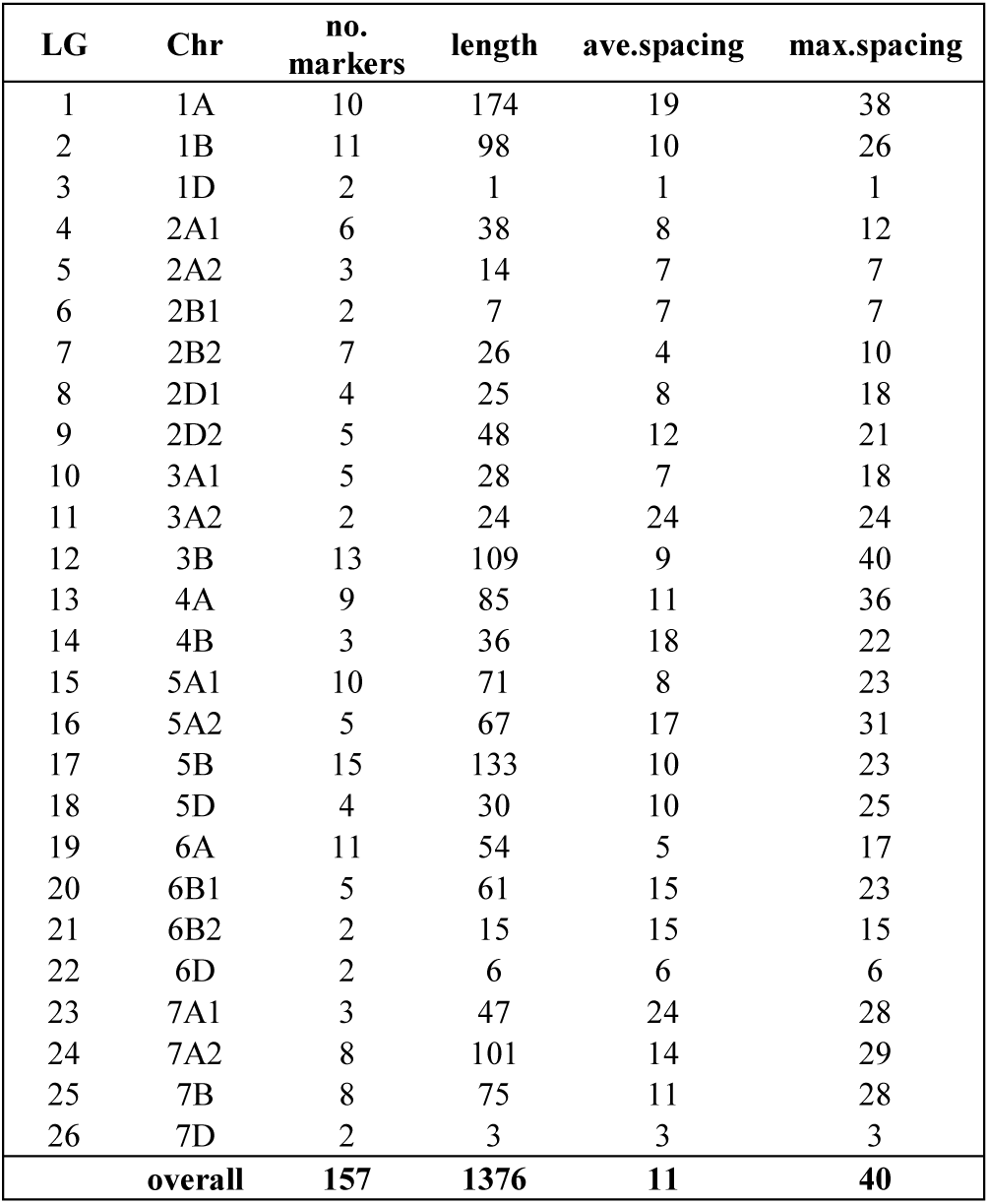
The table provides the number of markers, length, the average inter-marker spacing, and the maximum distance between markers, for each chromosome or linkage groups from the analysis. No markers have been mapped for 3D and 4D chromosome of wheat.

The threshold distance was set not higher than 30 cM between DNA markers along linkage groups (LG). Therefore, two LG have been obtained if there is a length of more than 30 cM between two neighboring DNA markers in pairwise analysis. In other words, if recombination fraction between two markers > 0.30%, more linkage groups formed.

### Basic statistics

General statistics provided in the paper followed the methodologies described in (Broman 2010) which is a gold standard for genetic map construction, using an open-source and powerful statistical environment, R (Ihaka and Gentleman 1996). The `gtl` library of the R-QTL package has been used to run whole analyses. In order to read the genetic dataset or genetic map saved as a CSV (Comma Separated) file (S.6), the `read.cross ()` function has been employed. Although the map was constructed using MapDisto and marker positions were known, `estimate.map ()` function has been set as `TRUE` in R, because the aim was to re-estimate genetic distances. If it is set to `FALSE,` marker distances along the chromosome would not be counted, and thus R draws the map with the same distances for all markers. During genetic map estimation, the probability of assumed genotyping error rate was set to 1e-4. The function `summary ()` provided the overall summary of the cross. To plot the homozygous-AA, homozygous-BB, heterozygous-AB alleles and missing genotypes as a heat map the `geno.image ()` function was used. In order to calculate either the numbers of genotyped/missing markers for each sample or the number of genotyped/missing samples for each marker the function `ntyped ()` has been used. The function `findDupMarkers ()` dedicated to find duplicate markers was applied to look only for markers that have exact matching genotypes and the same pattern of missing data. Before identifying duplicates, we used `comparegeno ()` function to count the proportion of matching genotypes between all pair of individuals so that we could evaluate how much the individuals are genetically close. To research the segregation ratio within the mapping population we utilized the function `geno.table ()` which forms a table containing information about samples/individuals with their genotypes at each molecular marker and calculates P-values for each molecular marker from chi-square tests for Mendelian segregation. A recombination fraction between all pairs of genetic markers was estimated by `est.rf ()` and was visualized using `plotRF ()` with minor amendments to the original usage of the function. In order to validate whether there are markers with recombination fractions > ½ and with high LOD scores, `pull.rf ()` function was used. Then obtained results was plotted using general `plot ()` function as numeric. The information on the number of markers, length, the average inter-marker spacing, and the maximum distance between markers, for each chromosome or linkage groups was generated by `summaryMap ().` The genetic map as a table with chromosome coordinates and marker names was obtained with a help of the `pull.map ()` function. To draw the genetic map, the `lmv.linkage.plot` function of the LinkageMapView package was used.

## Results

### Population development and genetic map construction

Here, we report the development of a segregating population derived from the cross of wheat varieties Paragon and Pamyati Azieva, specifically Recombinant Inbred Lines (RILs), and the construction of the first genetic map of RILs in collaboration between John Innes Centre (JIC), UK and Institute of Plant Biology and Biotechnology (IPBB), Kazakhstan. This is noteworthy, as this is the first mapping population developed and genetic map ever constructed, involving wheat cultivar adapted to the main wheat growing environments of Kazakhstan, using modern molecular marker technology. As a result, 94 Recombinant Inbred Lines have been obtained and the total length of constructed genetic map was 1376 cM. The total number of mapped SNP KASP markers was 157 which reside on 26 linkage groups. The largest linkage group was the linkage group 1 for 1A chromosome with 174 cM map size. By contrast, the linkage group 3 for 1D chromosome was the smallest in size with only 1 cM. The linkage group 17, with total of 133 cM map size and 23 cM of maximum spacing, was the densest, assigning 15 molecular markers (Table.1 and S.5). A total number of unassigned markers was 20 (not shown).

### DNA isolation

DNA was extracted from five- and six-days old seedlings grown in an incubator with a temperature range of 20-22°C. Regarding to Nanodrop outcomes (data is not provided) the quantity of isolated DNA was an average higher than 100 μl/ng and the quality varied between 1.8 – 2.0 (A260/A280) after purification with Qiagene Mini spin columns. As KASP requires less DNA concentration to set up PCR comparing to usual PCR, DNA of each recombinant line has been diluted separately with distilled water to make DNA with final concentration of 10 ng/μl.

### General statistics

The function `summary ()` showed that the type of a cross was “riself” as expected, with a meaning of RILs obtained by self-fertilization. As threshold distance between markers was set not to be higher than 30 cM, two linkage groups were obtained for chromosomes such as 2A, 2B, 2D, 3A, 5A, 6B and 7A. Therefore, total number of chromosomes was 26 and of genotyped markers was 157, excluding one of the two duplicate markers, BS00090234 and BS00022781, with identical marker genotypes. These duplicate markers located on 2B2 linkage group have been identified by `findDupMarkers` () function and one, especially BS00022781, have been omitted from further analysis using `drop.markers` (). Thus, constructed map contained 26 LGs namely 1A (with 10 SNP DNA markers), 1B (11), 1D (2), 2A1 (6), 2A2 (3), 2B1 (2) 2B2 (7), 2D1 (4), 2D2 (5), 3A1 (5), 3A2 (2), 3B (13), 4A (9), 4B (3), 5A1 (10) 5A2 (5), 5B (15), 5D (4), 6A (11), 6B1 (5), 6B2 (2), 6D (2), 7A1 (3), 7A2 (8), 7B (8) and 7D (2) (Table. 1).

To draw the genetic map with all alleles and missing genotypes as a heat map the `geno.image ()` function was used, but the data was treated as F2 generation. As a result of plotting data points, there was an acceptably low level of missing genotypes observed for both individuals and markers, therefore there was no reason to drop neither individuals nor markers (FIG.1). Note that one should read the data set in R as f2 generation in order to plot all alleles and missing data for inbred lines obtained by self-fertilization. If the data set is treated as self-fertilized inbred lines, the program plots heterozygotes as missing genotypes. Therefore, one could end up with more missing data in data set. The function `plotMissing ()` under R-QTL package could potentially be utilized to implement the same analysis, but it does not provide haplotype (allelic) information. Therefore, it plots only missing genotypes and thus, we would recommend to apply `geno.image ()`to provide a whole picture of the map.

**FIG 1.**
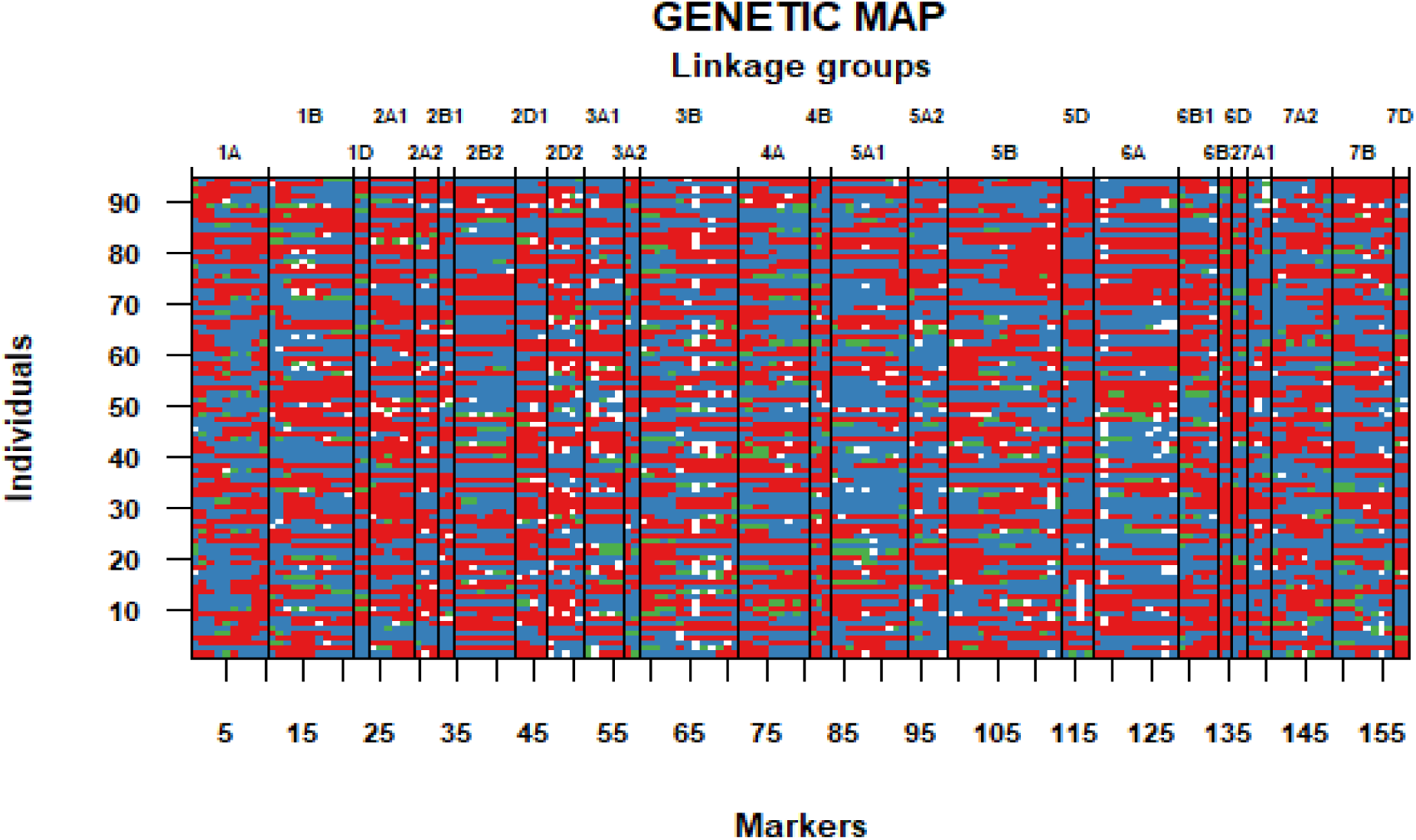
Genetic heat map: Red, blue and green refers to AA, BB and AB alleles respectively. White gaps are missing genotypes. Along on the top and bottom x axis linkage groups and markers are located respectively. Horizontal lines from top to down split linkage groups. The samples are potted on the y axis.

The genotyping success rate was 96.5% with 3.5% of missing data. Taken the percent genotyped as 100%, the share of AA and BB alleles would accounted for 47.7% and 47.2% respectively which is almost the same with remaining 5.1% of heterozygotes (AB) (FIG.2 and S.4). The plot of numbers of genotyped/missing markers for each individual and numbers of genotyped/missing samples for each maker using `ntyped ()` function has clearly shown that almost all of the individuals have been genotyped with more than 140 markers out of 157 with an exception of only three individuals, PamxPar-89, PamxPar-49 and PamxPar-48, which had 135, 136 and 137 genotyped markers respectively. There was only one marker, BS00060014, with more than 30 missing genotypes/samples out of 94 (FIG.3 and S.1). This confirms that we do not need to omit any individual and marker from analysis as those minor missing data (missing genotypes for both individual and marker) should not cause any difficulties.

**FIG 2.**
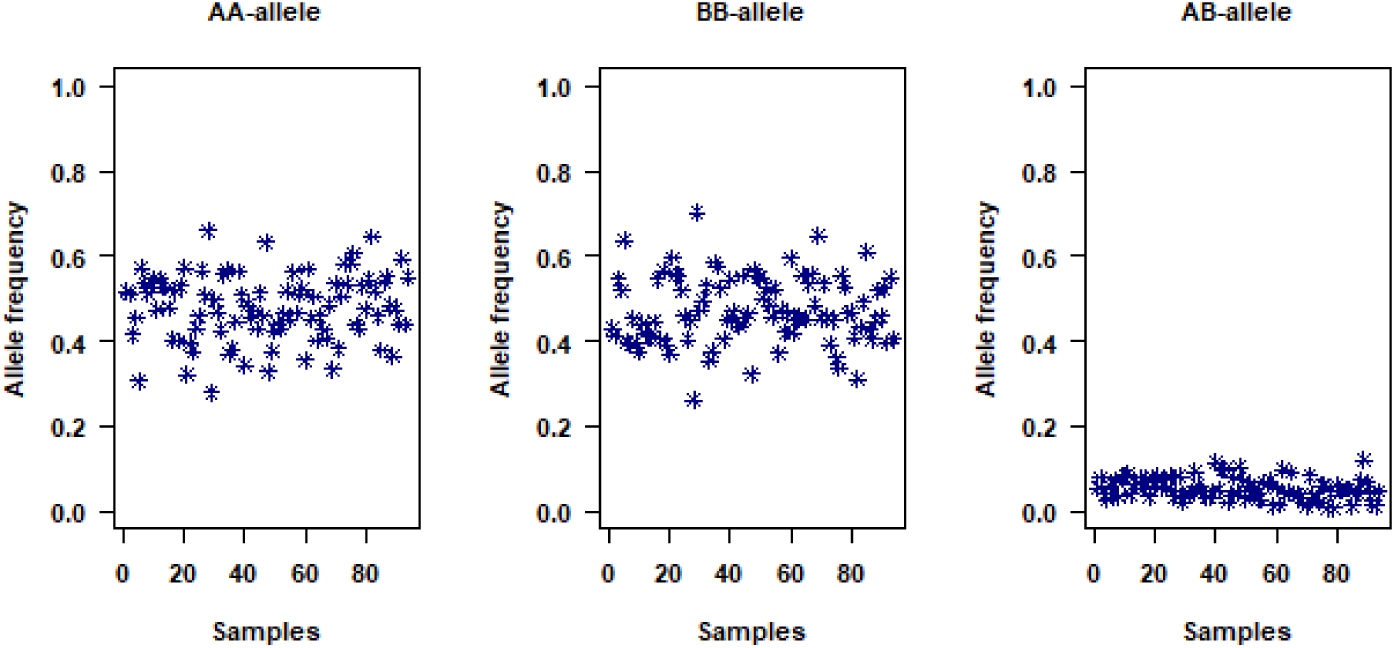
The proportion of AA and BB alleles were almost the same (47.7% and 47.2% respectively). By contrast AB genotypes consisted of only 5.1%, as expected in F4, suggesting that the population was mostly fixed at the majority of markers.

As a result of `comparegeno` function, we saw that Inbred Lines derived from the cross of PamxPar share about ∼45% genotypes at markers. We also could see individuals with highly similar genotypes, with more than 90%. Additionally, those individuals with almost similar genotypes, RIL30 with RIL31 (share 92.3% genotypes), RIL58 with RIL61 (90.1%) and RIL41 with RIL62 (90.2%), have been identified (FIG.4 and S.2).

**FIG 3.**
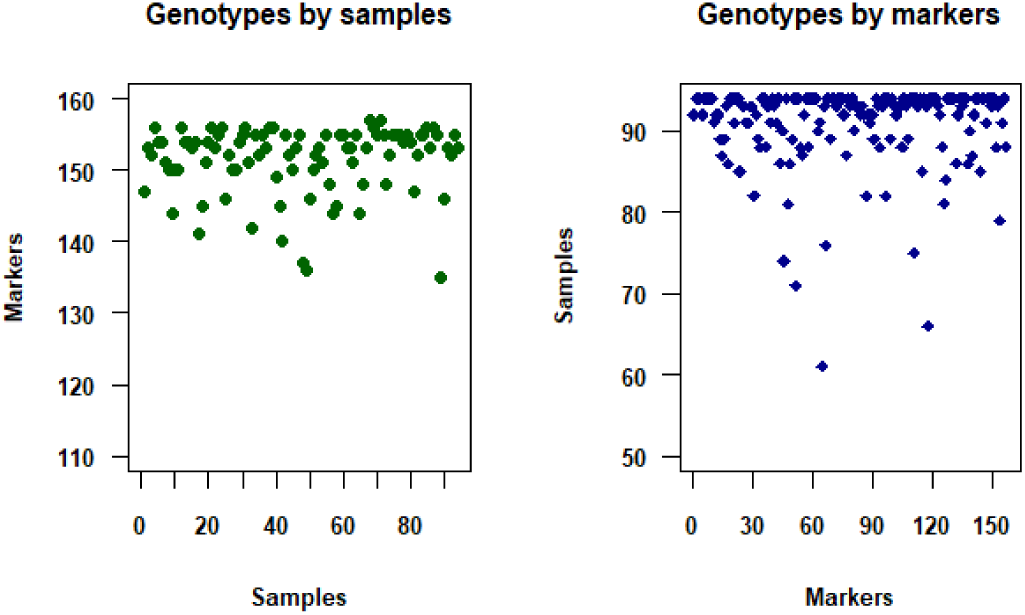
The number of genotyped markers for each sample (left) and the number of genotyped individuals for each marker (right).

**FIG 4.**
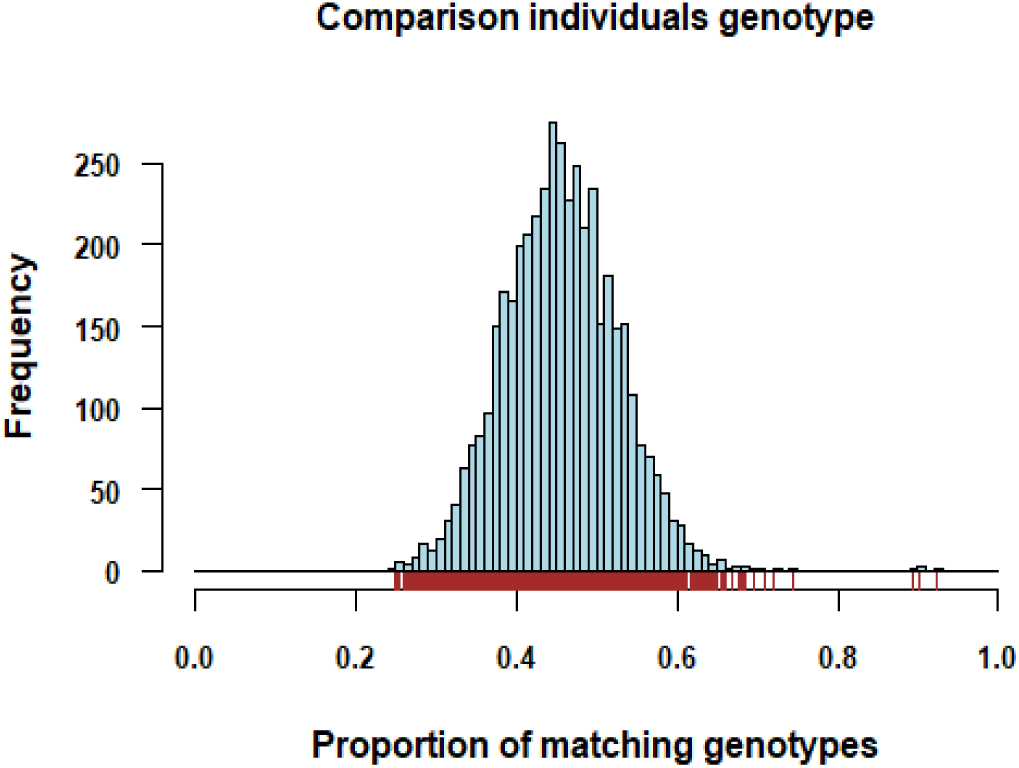
Histogram shows the comparison of genotypes for each pair of individuals.

The function `geno.table ()`provided the number of AA, BB, AB and missing genotypes with P-value obtained from chi-square tests of Mendelian segregation. As a result, only two markers, BS00085589 and BS00110651, located on 6B2 linkage group, showed slight segregation distortion towards AA allele, but P-value was lower than 0.05% (S.3).

As a result of the estimation of pairwise recombination frequency of all pairs of genetic markers, likely translocations between 2A1:2D1, 2A1:4A and 5A1:4A, was determined in the mapping population. This was identified when the recombination fractions for all pairs of markers and the LOD scores for tests of linkage between pairs of markers have been plotted as a grid (FIG.5).

**FIG 5.**
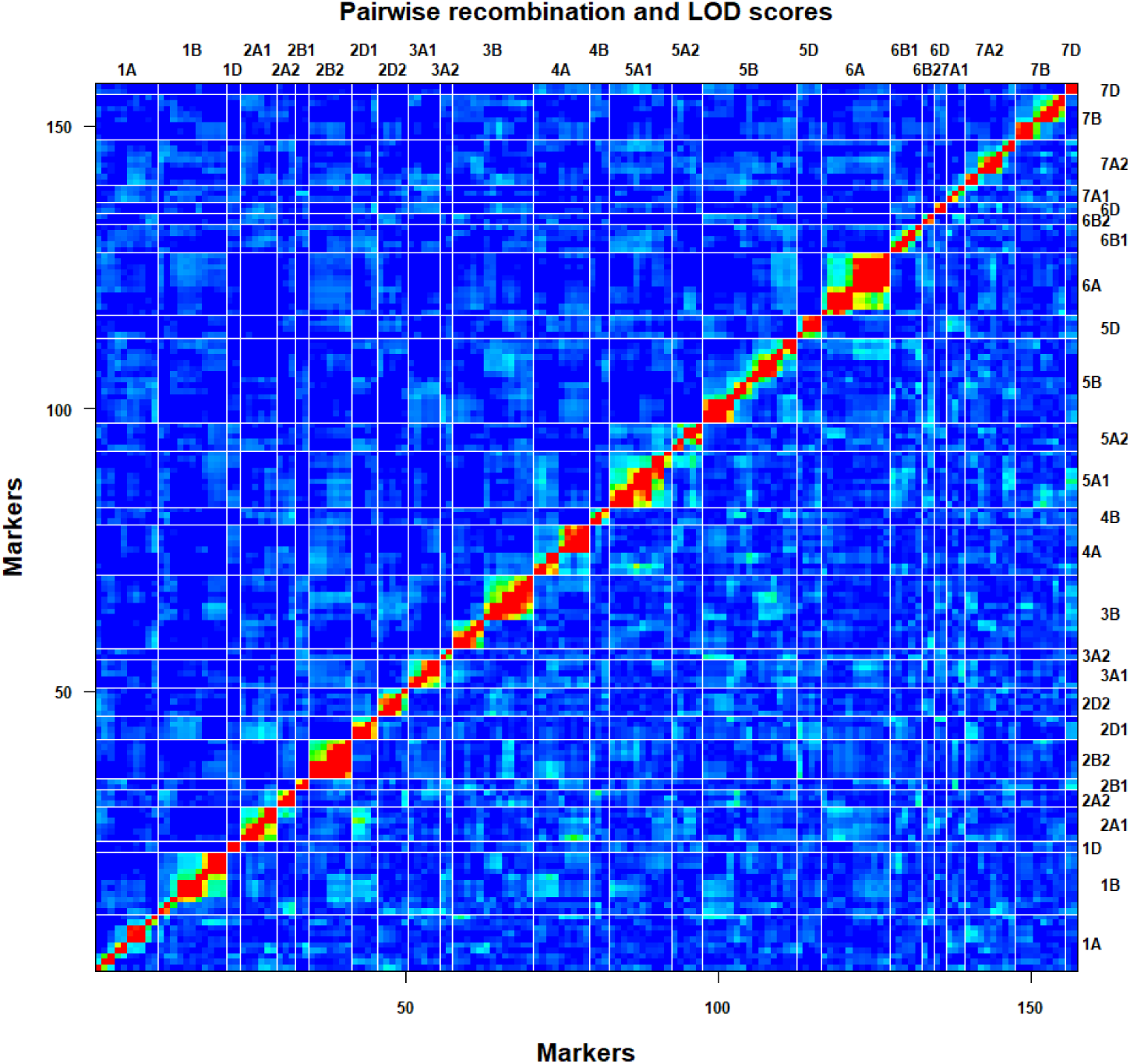
Pairwise recombination frequency and linkage analysis between markers. The recombination fractions are in the upper left triangle while the LOD scores are in the lower right triangle. Red corresponds to a large LOD or a small recombination fraction. In comparison blue represents a small LOD score and a high recombination fraction. A light grey indicates missing values.

Marker BS00078489 on LG 2A1 has been shown to be linked with three markers, BS00043985, BS00022730, BS00022276 and BS00097249, BS00022771, BS00060226, on LG 2D1 and LG 4A respectively. Moreover, SNP marker BS00065607 on LG 5A1 was linked with BS00022987 on LG 4A. However, when we looked at the real quantitative data, the LOD score of all marker pairs was too low. Recombination frequency was < ½ for 2A1: 2D1 and 5A1: 4A, but > ½ for 2A1: 4A (FIG.6 and Table 2). While no cytological evidence is available to support the outcome, we concluded no major translocations have occurred within biparental mapping population.

**FIG 6.**
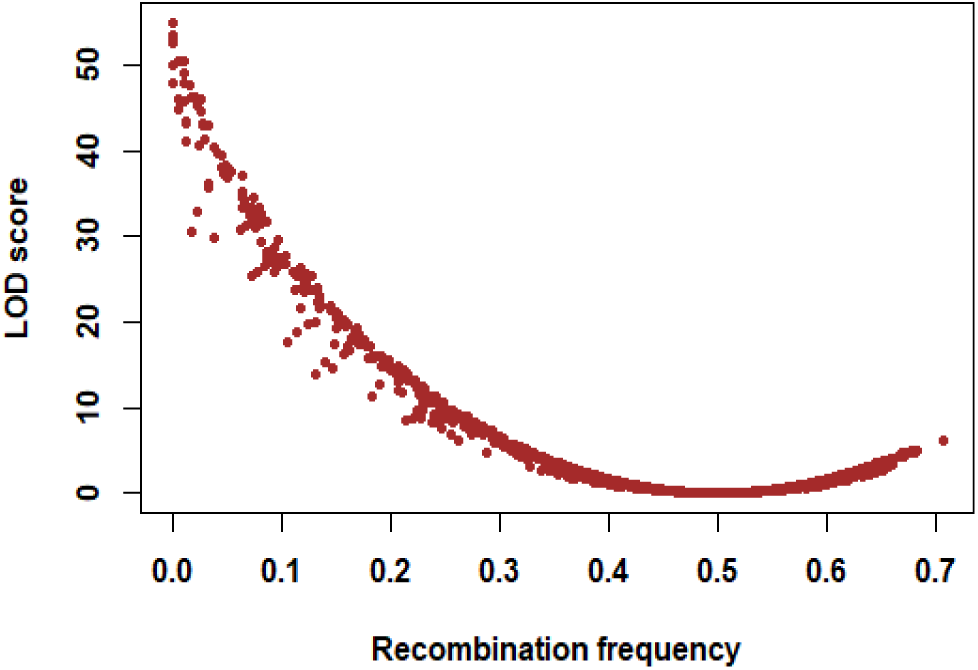
Plot of calculated LOD scores versus estimated recombination fractions as numeric for all marker pairs from est.rf () showed that some markers had RF more than ½, but LOD was low.

**Table 2.**
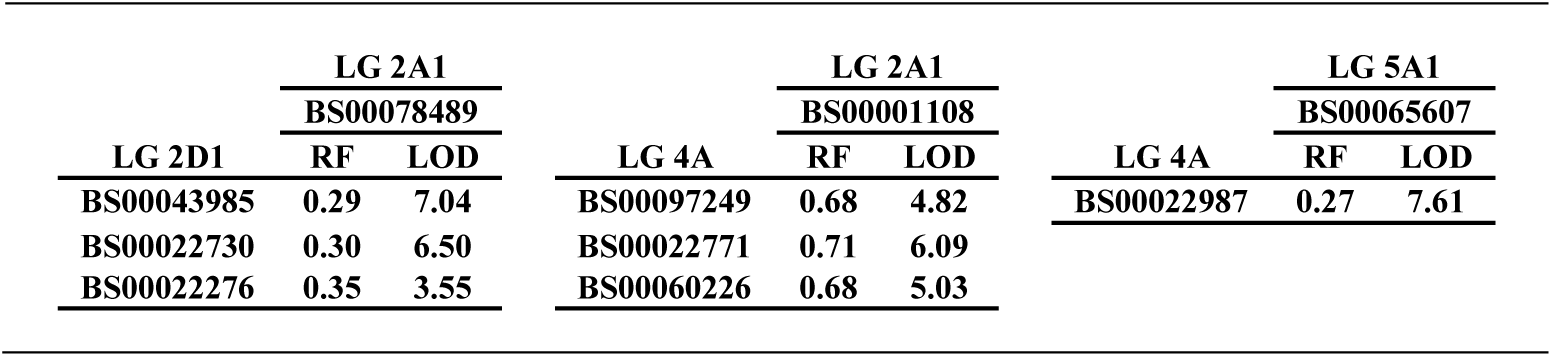
The table provides quantitative data for likely translocations determined based on the plot of LOD and RF. LG, RF and LOD stands for linkage groups, recombination frequency and LOD scores respectively.

## Conclusion

As the development of mapping population and construction of its genetic map is the essential initial step of identification of genes, associated with important plant traits, and the introgression of those identified genes into elite wheat varieties to optimise final grain harvest, the importance of this study is crucial. Even though most part of the research, including hybridization experiments and development of a mapping population, conducted using the facilities of the John Innes Centre, UK, there is a great potential to transfer this essential acquired knowledge and experience to Kazakhstan, if there is an interest from Kazakh government or private sectors to build such facilities in the country. Implementing so, would enable the Kazakh government to use country’s enormous agricultural capacity. This in turn, would optimise as well as centralise country’s plant and animal breeding programs both of which are not in satisfactory condition currently. The support of this kind of pre-breeding research would allow to strengthen the international collaborations which results in designing successful breeding programs. Putting plant and animal breeding programs into right path of development in Kazakhstan would contribute to country’s as well as world’s future food security.

## Supporting information

Genotypes by marker and individual

The pairwise comparison of genotypes for all pairs of individuals

Segregation ratio

Allele frequency

Genetic map

Genotyping dataset

## Supplementary Material

S.1: Genotypes by marker and individual.

S.2: The pairwise comparison of genotypes for all pairs of individuals.

S.3: Segregation ratio.

S.4: Allele frequency.

S.5: Genetic map

S.6: Genotyping dataset

## Acknowledgments

We really thank funding bodies: ADAPTAWHEAT (289842) funded by EU 7th Framework Programme for Research and Technological Development and the grant 1784/ΓΦ4 funded by Ministry of Science and Education of Republic of Kazakhstan, and “BOLASHAK” International Presidential Scholarship.

## Contributions

SG and YT initiated and designed the study. KY was involved in genotyping and conducted statistical analysis and re-estimated the genetic map using R and wrote the article. SG reviewed and edited the article. SO, M Ch and ML developed the mapping population and conducted genotyping and estimated initial genetic linkage groups using MapDisto. AA conducted field experiments on RILs in Kazakhstan. SA supplied the plant material and supervised the research in Kazakhstan

