## Supplementary figures and images for "Constrution of a genetic map of rils derived from wheat (*T. aestivum* L.) varieties pamyatii azieva x paragon using high-throughput snp genotyping platform kasp - *competative allele specific PCR*"

### Genetic map

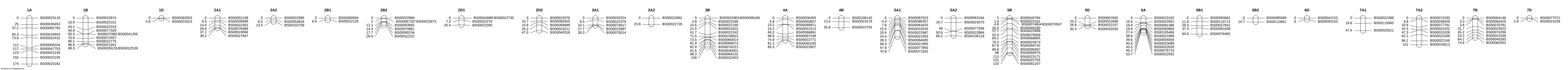
